# Light remote control of alternative splicing in roots through TOR kinase

**DOI:** 10.1101/472126

**Authors:** Stefan Riegler, Lucas Servi, Armin Fuchs, Micaela A. Godoy Herz, María G. Kubaczka, Peter Venhuizen, Alois Schweighofer, Craig Simpson, John W.S. Brown, Christian Meyer, Maria Kalyna, Andrea Barta, Ezequiel Petrillo

## Abstract

For plants, light is the source of energy and the most relevant regulator of growth and adaptations to the environment by inducing changes in gene expression at various levels, including alternative splicing. Chloroplasts trigger retrograde signals that control alternative splicing in leaves and roots in response to light. Here we provide evidence suggesting that sugars, derived from photosynthesis, act as mobile signals controlling alternative splicing in roots. The inhibition of TOR kinase activity diminishes the alternative splicing response to light and/or sugars in roots, showing the relevance of the TOR pathway in this signaling mechanism. Furthermore, disrupting the function of the mitochondria abolishes alternative splicing changes supporting a key role for these organelles in this signaling axis. We conclude that sugars can act as mobile signals coordinating alternative splicing responses to light throughout the whole plant, exerting this function in roots by activating the TOR pathway.

**Graphical Abstract:** 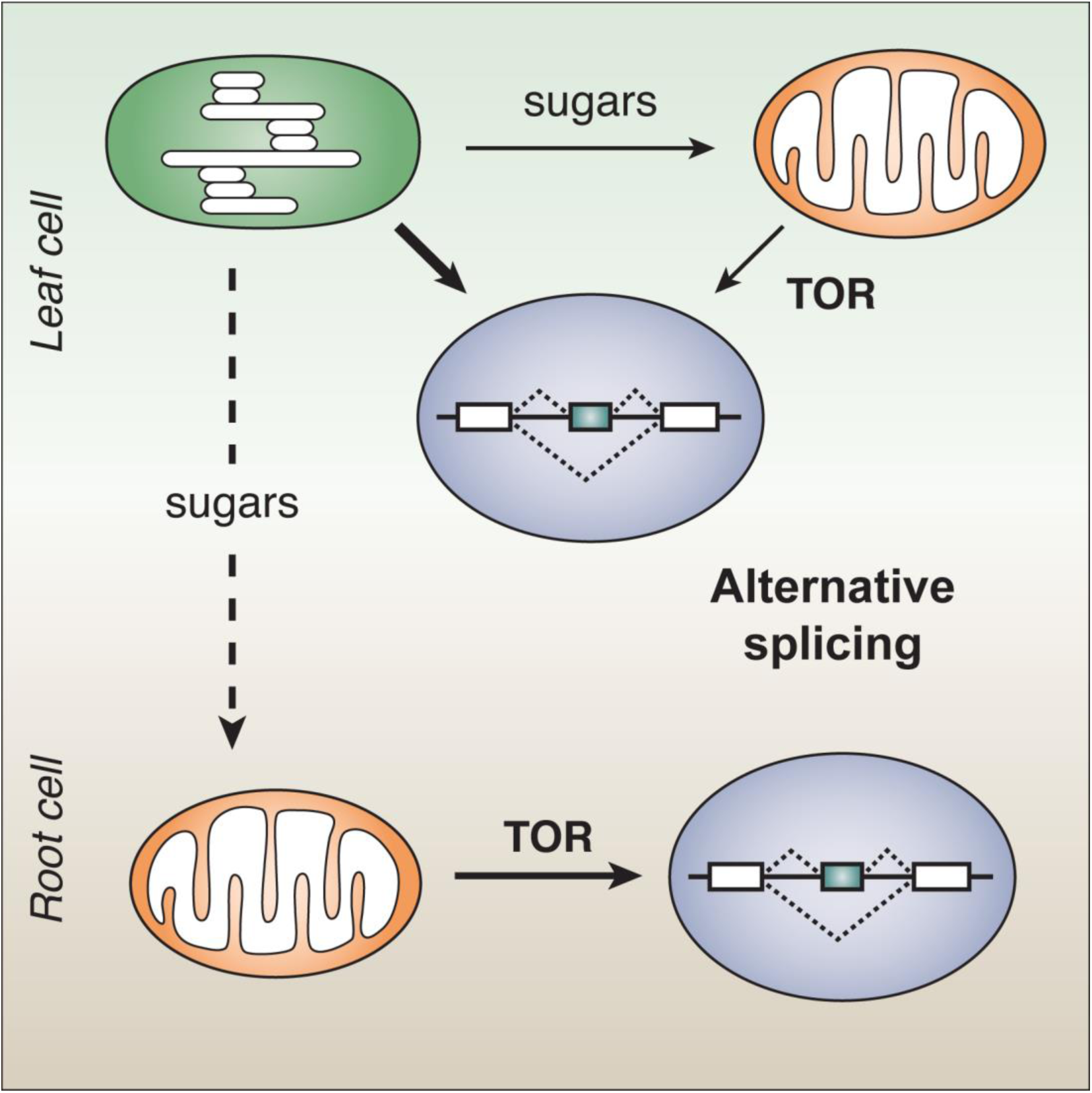

Art by Dr. Luciana Giono.

## Introduction

Light is essential for plants, it is their source of energy and makes carbon fixation possible allowing life on Earth as we know it, while it is also the main source of information about the ever-changing environment. Since plants’ success relies on this environmental cue, it is not surprising that in the course of evolution plants have developed a rich variety of means to sense light wavelength, direction and quantity (1, 2). Different families of photo-sensory proteins are important for proper light perception. However, chloroplasts – the organelles responsible for photosynthesis - are nowadays gaining relevance as key components of plant photo-sensory networks (3, 4). The nucleus controls most aspects of organelle biogenesis and function by means of anterograde signals. Conversely, chloroplasts and mitochondria regulate plant gene expression with retrograde signals that modulate transcription and translation of nuclear genes (5, 6). Previously, we reported that nuclear alternative splicing is modulated by light through a chloroplast retrograde signaling pathway (7–9). Interestingly, photosynthesis also modulates alternative splicing responses in the roots, suggesting the existence of intra- and inter-cellular mobile retrograde signals. The central question that now arises is how light coordinates nuclear splicing throughout the whole plant.

Mature chloroplasts generate carbon compounds that are further metabolized in the very same leaves or loaded into the phloem to feed distant non-photosynthetic tissues (e.g., roots). Sucrose is the main sugar loaded into the phloem travelling from green-productive-source tissues to sinks. As in the case of light, sugars have a dual role in plant cells serving as sources of energy and signals (10). Sugar levels modulate gene expression, metabolism, cell cycle, development and responses and adaptation to the environment (11, 12). Different intra- and extra-cellular sensors can perceive sugars. In addition to these direct pathways, where the hexokinase 1 (HXK1, AT4G29130) has a central role, sugars can also be indirectly perceived by the sucrose non-fermenting related protein kinase 1 (SnRK1) and by the target of rapamycin (TOR) kinase. Sugars repress SnRK1.1 (AT3G01090) (13) and activate TOR kinase (AT1G50030) (14). Sugars derived from photosynthesis can move and activate the meristem in the root (15) and modulate G protein activation in a long-distance communication pathway (16). Furthermore, these molecules can regulate alternative splicing in etiolated seedlings in a manner similar to light (17). Here, we show that by activating the TOR pathway in the roots, sugars could act as a mobile signal, coordinating alternative splicing responses to light throughout the whole plant.

## Results

### Sucrose changes alternative splicing mimicking light

Light regulates alternative splicing of the serine/arginine-rich splicing factor, *At-RS31* (AT3G61860, Fig. 1A-B) in *Arabidopsis thaliana*. The effects of light on this event were shown by different groups working with plants incubated in different conditions (4, 7, 18), making it a robust endogenous reporter for splicing regulation by this environmental cue. Previously we showed that chloroplasts act as remote controls of alternative splicing, reaching root cell nuclei with their signals (7). Sugar addition to the plant growth media (see Suppl. Fig. 1 and methods for details), though not eliminating *At-RS31* alternative splicing responses to light/dark in seedlings, does reduce the splicing index (SI = ratio of the longest isoform to all the isoforms of a gene), resembling the action of light (Fig. 1B and (7)). Moreover, in other alternative splicing events, i.e., the translation initiation factor 1A, eIF-1A (AT5G3568), and the SNF1-RELATED PROTEIN KINASE 1.2, SNRK1.2 (AT3G29160), the addition of this sugar completely phenocopies light (Suppl. Fig. 2). These findings, together with a recent report showing that sugars modulate alternative splicing in etiolated seedlings in a similar manner as light (17), prompted us to hypothesize that sugars, produced in photosynthetic cells, control alternative splicing in non-photosynthetic tissues. As explained above, when light/dark treatments are performed on whole plants (Fig. 1C), light regulates *At-RS31* alternative splicing in leaves (Fig. 1E) and roots (Fig. F). However, when plants are dissected before the light/dark treatment (Fig. 1D), roots disconnected from photosynthetic tissues lose the capacity to change the alternative splicing of *At-RS31* in response to light (Fig. 1F and (7)). In this treatment scheme, sucrose addition completely recapitulates light effects in roots, detached or not (Fig. 1F), with milder influence in leaves (Fig. 1E). These results demonstrate that sucrose modulates nuclear alternative splicing of *At-RS31* in root cells. The difference between the effect of sucrose in leaves and roots may be due to an inefficient sugar uptake by leaves from the agar-growth media. We applied vacuum infiltration to increase the uptake of these compounds in the plants flooded with liquid growth media (with sucrose or sorbitol as control). In these conditions, sucrose mimics light effects on the alternative splicing patterns of *At-RS31*, *At-U2AF65A* (AT4G36690) and *At-SR30* (AT1G09140), in leaves as it does in roots (Figs. 1G-H and Suppl. Fig. 3). Furthermore, this is independent of the chloroplasts, as shown by the fact that blocking the photosynthetic electron transport with DCMU (3-(3,4-dichlorophenyl)-1,1-dimethylurea (19)) does not disrupt sucrose effects on these events (Fig. 1G-H, Suppl. Fig. 3).

**Figure 1.**
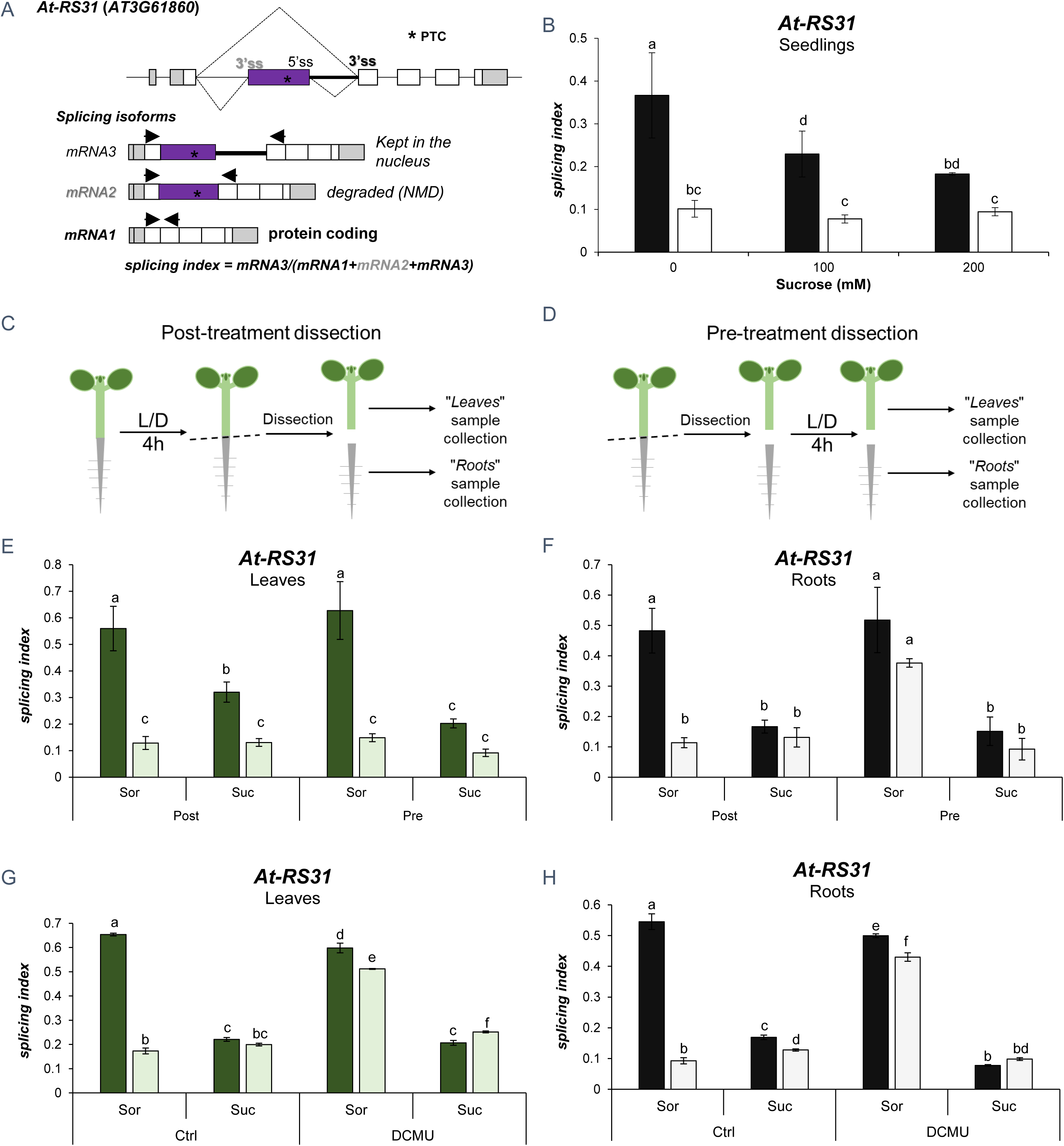
Sucrose changes alternative splicing in the same manner as light. **A**) Gene model and alternative splicing isoforms of *At-RS31.* *, PTC: premature termination codon. Arrow heads: primers for splicing evaluation. **B**) Sucrose (Suc) affects alternative splicing in plants. Sucrose addition to plant growth media at 100 mM or 200 mM concentration produces a shift in the splicing index (SI) values of *At-RS31* similar to that triggered by light. Sorbitol (Sor) was used as osmotic control at 200mM or together with sucrose at 100 mM to reach the same osmolarity in all treatments. **C-F**) Sucrose mimics the effects of light in roots. *Arabidopsis thaliana* (Col-0) plants were incubated under a light/dark protocol for about four hours (Suppl. Figure S1). Incubation with sugar was done during the light/dark (L/D) treatments. Post-treatment dissection (**C**) was done after the light/dark treatment, immediately before sample collection. Pre-treatment dissection (**D**) was done prior to light/dark incubation. After 48 hours of darkness, transferring the plants to light results in a rapid change in the SI of *At-RS31*. Sucrose addition (200 mM) causes a small shift of SI in leaves, detached or not (**E**), while fully mimics the effect of light in roots (**F**). Sorbitol 200 mM was used as an osmotic control. **G-H**) Sugar effects on alternative splicing do not involve the chloroplasts. Increasing sucrose (100 mM) uptake by leaves using vacuum infiltration (5 min) generates similar effects of sugars in the different tissues. Blocking chloroplast electron transport with DCMU (20 µM) abolishes light effects in leaves and roots without affecting sugar induced changes. Ethanol was used as control (Ctrl). Incubations with drugs started 1 h before light/dark treatments and were maintained until sample collection. Plants were grown for two weeks in constant light (~100 µmol/m2s). See Suppl. Fig. 1 for more details. **B, E-H**) Lighter bars, light; darker bars, darkness. The graphs show splicing index means ± standard error (n=4). Same letters indicate means that are not statistically different (p>0.05).

**Figure 2.**
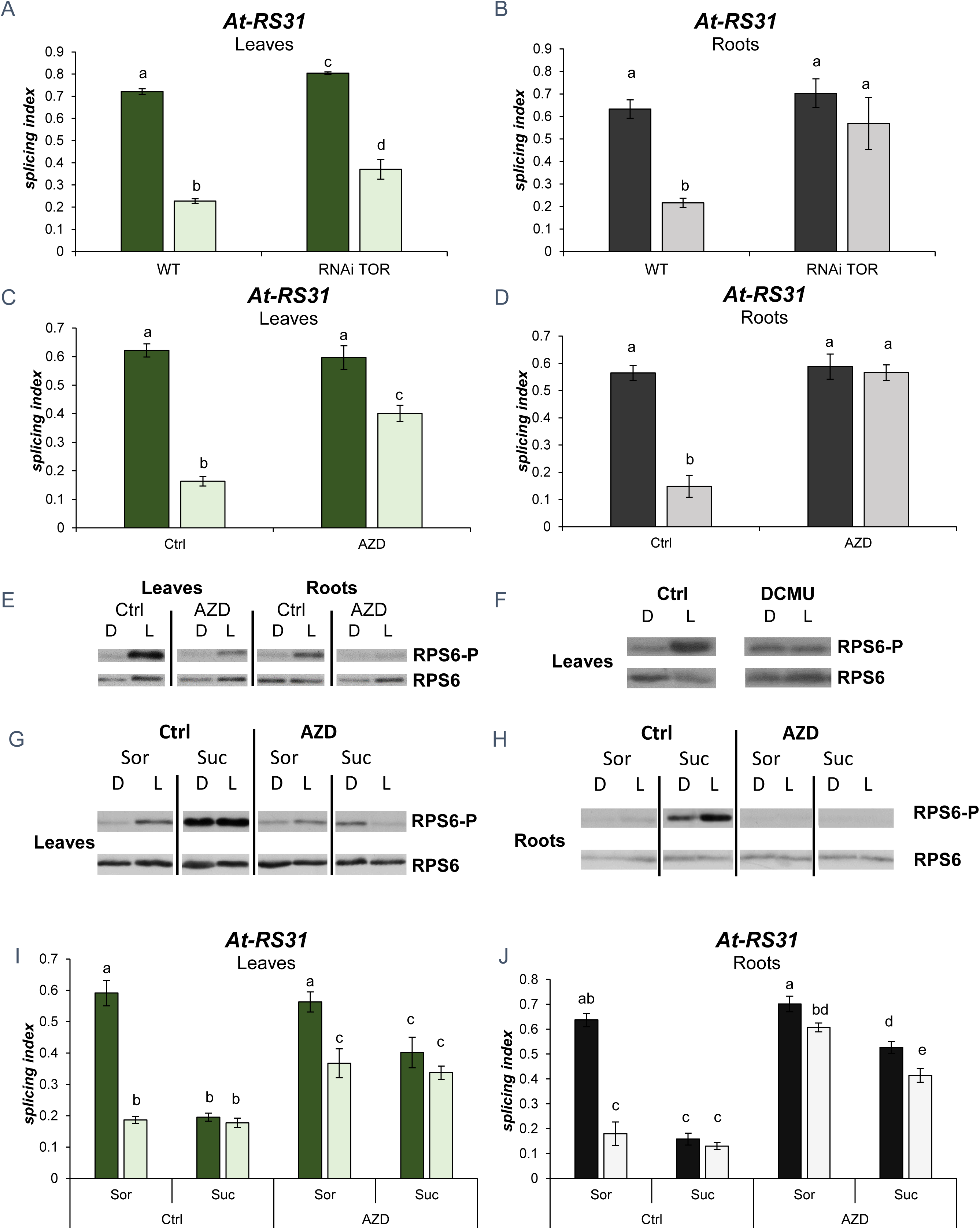
TOR kinase modulates alternative splicing responses to light and sugars. **A-B**) TOR knockdown disrupts *At-RS31* alternative splicing responses to light. Light/dark-induced *At-RS31* alternative splicing changes are diminished in leaves (**A**) while completely abolished in roots (**B**) of an RNAi Tor line (35-7 TOR RNAi line (20)). *Col*, *Col-8*. **C-E**) TOR kinase is necessary for light-mediated alternative splicing response in roots. TOR kinase activity inhibition by AZD-8055 (AZD) diminishes light effects on *At-RS31* alternative splicing in leaves (**C**) while completely abolishes the changes in roots (**D**). Ctrl, dimethyl sulfoxide (control). **E**) Ribosomal protein S6 (RPS6) phosphorylation (-P) is increased by light (L) in leaves and roots in comparison to dark (D). AZD-8055 (AZD) abolishes the changes in RPS6 phosphorylation induced by light. **F**) Blocking the photosynthetic electron transport inhibits the phosphorylation of RPS6. DCMU (15 µM) blocks the electron transport in the chloroplasts and the light-induced phosphorylation of RPS6 in leaves. **G-H**) Sucrose addition greatly increases RPS6 phosphorylation in leaves (**G**) and roots (**H**) but this effect, as the effect of light, is inhibited by AZD (20 µM). RPS6-P, phosphorylated RPS6. RPS6, total RPS6. **I-J**) TOR inhibition abolishes light and sugar triggered alternative splicing changes. Leaf responses (**I**) to light and sugars are diminished by AZD while root responses (**J**) are completely abolished. Plants (*Col-0*) were flooded with liquid MS-MES supplied with sucrose (Suc) 100 mM or sorbitol (Sor) 100 mM as an osmotic control during the light/dark treatment. Light/dark treatments were done as described for post-treatment dissection in Figure 1. Incubations with drugs started 1 h before light/dark treatments and were maintained until sample collection. Vacuum infiltration was used to increase different compounds’ uptake. **A-D, I-J**) Lighter bars, light; darker bars, darkness. The graphs show splicing index means ± standard error (n=4). Same letters indicate means that are not statistically different (p>0.05).

**Figure 3.**
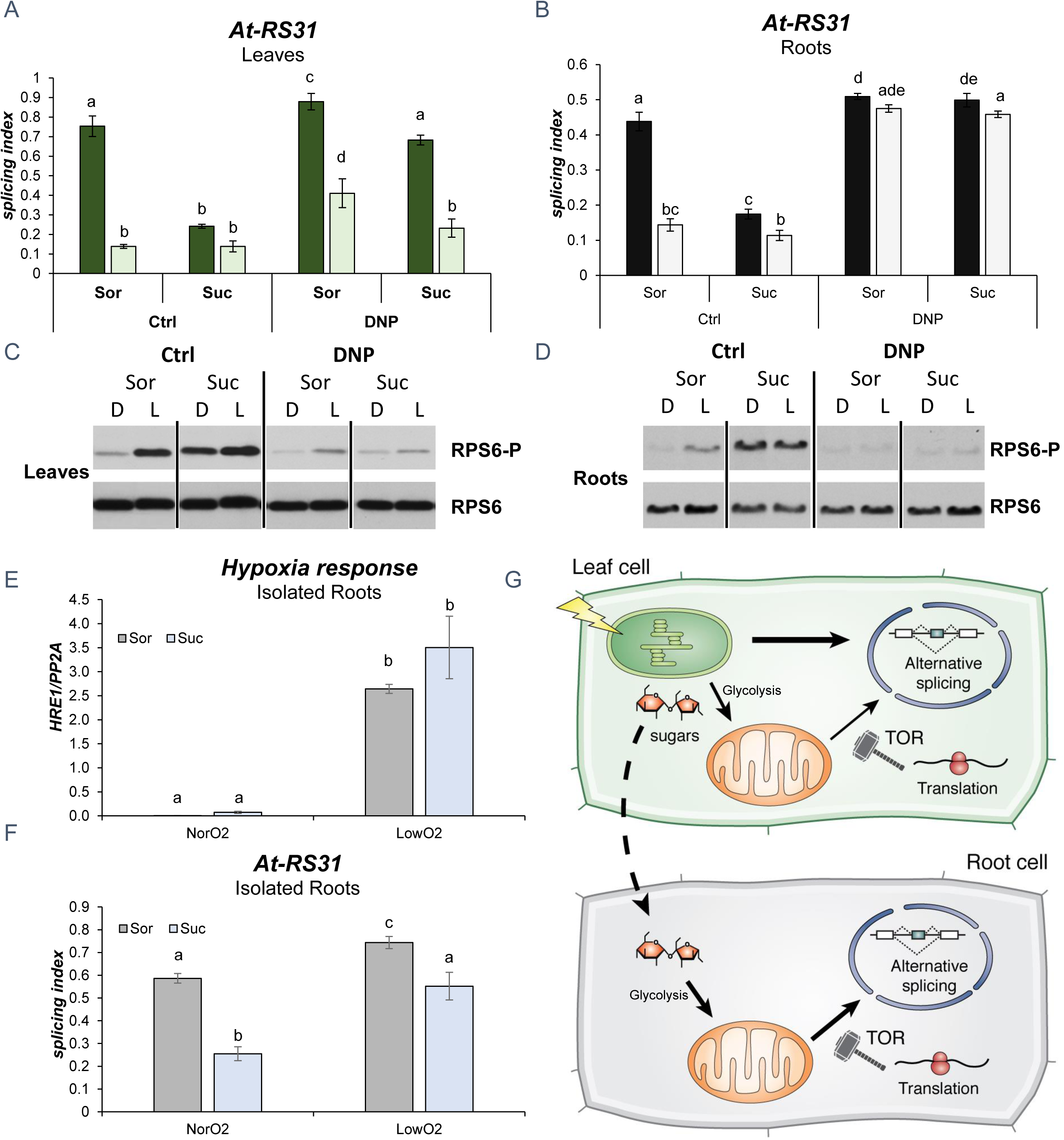
Light and sugar signaling in roots relies on mitochondrial activity. **A-D**) The ionophore 2,4-dinitrophenol (DNP) that causes proton gradient dissipation across membranes, is not abolishing light-induced changes in *At-RS31* splicing in leaves (**A**), however, sucrose (Suc) effects are diminished by it. Alternative splicing control in roots (**B**) is fully dependent on mitochondrial activity since DNP treatment completely abolishes light and sucrose-induced splicing index changes. Concomitantly, DNP treatment abolishes light and sucrose-induced phosphorylation of RPS6 in leaves (**C**) and roots (**D**). **A-B**) Lighter bars, light; darker bars, darkness. The graphs show splicing index means ± standard error (n=4). Same letters indicate means that are not statistically different (p>0.05). **A-D**) Plants were grown in constant light, transferred to darkness for 48 hours and then treated with DNP (20 µM) and sucrose (Suc) 100 mM, or sorbitol (Sor) 100 mM as osmotic control, for further 4 hours in light or dark (see Suppl. Fig. 1 for more details). Dimethyl sulfoxide was vehicle/control (Ctrl) treatment for DNP. **E**-**F**) Mitochondrial respiration is relevant for isolated roots to change alternative splicing in response to sucrose. **E)** *HRE1* (*hypoxia-inducible ethylene response factor 1*) expression is increased in low oxygen (LowO2) in comparison to air (NorO2). **F)** An atmosphere with reduced oxygen levels, replaced by nitrogen saturated air, attenuates *At-RS31* alternative splicing changes in response to sucrose in isolated roots. **E-F**) Plants were grown in constant light for two weeks, transferred to darkness for 48 hours (see Suppl. Fig. 1). Seedlings were dissected, and isolated roots treated with 50 mM sucrose (Suc) or sorbitol (Sor) as a control. The graphs show splicing index means ± standard error (n=3). Same letters indicate means that are not statistically different (p>0.05). Statistical analyses were done using InfoStat with Fisher LSD for comparisons. **G**) Model: different retrograde signals regulate alternative splicing through inter-organellar communication. Photosynthesis in the chloroplasts generates different signals that activate translation and modulate alternative splicing in the nucleus of photosynthetic leaf cells. Among these signals, sucrose, which is transported to the roots and feed the mitochondria after glycolysis, activates TOR kinase and the translation machinery, thus changing the alternative splicing in this heterotrophic tissue.

### TOR kinase modulates alternative splicing responses to light and sugars

Sugars derived from photosynthesis are known to modulate the activity of different plant kinases, like HXK1, SnRK1 and TOR kinase (12). In our previous report (7), we ruled out HXK1 and SnRK1.1 (AT3G01090) as relevant sensors for the signaling pathway affecting alternative splicing responses to light in seedlings. Hence, we evaluated the involvement of TOR kinase, another sugar/energy sensor. Light/dark treated TOR RNAi transgenic plants show a small reduction in alternative splicing responses in leaves (Fig. 2A), evidencing a minor contribution of the TOR pathway in this tissue. In roots, however, the TOR knockdown line shows a complete abolishment of the light effects on the alternative splicing of *At-RS31* (Fig. 2B). Similar results were obtained with *At-U2AF65A* and, partially, with *At-SR30* (Suppl. Fig. 4), suggesting that TOR kinase is a key factor for the alternative splicing regulation by light in root cells. Nevertheless, since low levels of TOR kinase protein produce extremely short roots (20), we decided to use a specific inhibitor to discard an indirect effect of the RNAi lines. AZD-8055 (AZD), an ATP competitive TOR inhibitor (21), diminishes light-induced changes in the splicing index of *At-RS31* in leaves (Fig. 2C), while completely abolishing these changes in roots (Fig. 2D), exactly in the same way as observed with the TOR RNAi line. These results point to a key and direct role of TOR kinase activity in the light/sugar axis modulating alternative splicing. This conclusion is further supported by the fact that AZD disrupts sucrose-induced changes in the alternative splicing of *At-RS31*, *At-U2AF65A* and *At-SR30,* in isolated (detached) roots, which are not directly responsive to light but responsive to sugars (Suppl. Fig. 5 and Fig. 1F).

Finding a role for TOR kinase in this signaling pathway prompted us to evaluate the activity of the TOR signaling pathway in light/dark and under different treatments, to correlate it with the splicing changes. We used the ribosomal protein subunit 6 (RPS6) phosphorylation as a proxy for TOR pathway activity (22). Figure 2E shows that light activates the TOR signaling pathway in leaves and roots and that AZD efficiently inhibits this activation in both tissues. Similarly, this activation by light is abolished in the TOR-RNAi line (Suppl. Fig. 6). Since the signaling pathway that controls *At-RS31* alternative splicing involves the chloroplast, we asked if blocking the photosynthetic electron transport would abolish TOR activation triggered by light. The treatment with DCMU completely inhibits RPS6 phosphorylation induced by light in leaves (Fig. 2F), suggesting that, indeed, chloroplast function, TOR kinase activity and alternative splicing regulation are strongly linked. Furthermore, AZD treatment disrupts sucrose effects in roots and leaves, both on the phosphorylation of RPS6 (Fig. 2G-H) and, to different degrees, on the alternative splicing of *At-RS31* (Fig. 2I-J), *At-U2AF65A* and *At-SR30* (Suppl. Fig. 7). Importantly, the inhibition of RPS6 phosphorylation induced by sucrose in both tissues rules out a difference in the uptake and/or efficacy of AZD between leaves and roots. These results indicate that a light/sugar axis is of key relevance for alternative splicing regulation in roots, mainly, through the activation of the TOR kinase pathway.

### Light and sugar signaling in roots relies on mitochondrial activity

Xiong and colleagues have shown that the activation of the TOR pathway in roots occurs via sugars derived from photosynthesis that, after glycolysis in this tissue, feed the mitochondria. This signaling pathway affects root gene expression and activates the meristems (15). At-RS31 protein accumulation in roots occurs, mainly, in the meristems (Suppl. Fig. 8). Its abundance correlates with the splicing pattern in the different conditions. The higher the abundance of the *mRNA1* isoform (the coding transcript), the higher the protein levels, as occurs in light vs. dark conditions (Suppl. Fig. 9). We hypothesized that a pathway similar to the one activating root meristems via mitochondria and TOR kinase could be responsible for the regulation of *At-RS31* alternative splicing by light/sugars in roots. Following this line of reasoning, uncoupling the energy generation from the electron transport in the organelles using an ionophore (2,4-dinitrophenol, DNP) should abolish the alternative splicing changes induced by light and sucrose. In leaves, this uncoupler has only a mild effect on light-mediated *At-RS31* alternative splicing changes, while it clearly reduces sucrose effects restoring light/dark responses in this tissue (Fig. 3A). In roots, however, DNP completely blocks the changes in *At-RS31* alternative splicing induced by light and/or sucrose (Fig. 3B). These results suggest that mitochondria have a key role in the light/sugar signaling pathway in roots. Interestingly, the addition of DNP increases the light splicing indexes of *At-RS31* (Fig. 3A), *At-U2AF65A* and *At-SR30* in leaves (Suppl. Fig. 10), mirroring the effects of AZD and the TOR RNAi knockdown. This observation prompted us to analyze whether DNP-triggered changes were related to the activity of the TOR pathway. Remarkably, this uncoupler inhibits the phosphorylation of RPS6, i.e., the activation of the TOR pathway, induced by light and/or sucrose in leaves and roots (Fig. 3C-D), once more, resembling the effects of AZD and TOR knockdown. In conclusion, DNP abolishes TOR activation by sugars in leaves and roots. It is, however, important to mention a small increase in the phosphorylation of RPS6 in light-treated leaves that is resistant to TOR inhibition and the disruption of proton gradients across membranes (Figs. 2G, 3C and Suppl. Figs. 6 and 10). Furthermore, these results were independently validated following an inhibitor-independent approach. We generated a low oxygen atmosphere, that was evidenced by the expression of *HRE1,* a hypoxia-responsive gene (23) (Fig. 3E). Under these conditions, isolated roots show reduced *At-RS31* alternative splicing changes in response to sucrose (Fig. 3F). Taken together, these results indicate that alternative splicing regulation of different genes in root cells is connected to the activity of the mitochondria and the concomitant activation of the TOR kinase pathway. As a corollary, the fact that the alternative splicing changes triggered by light in leaves are not abolished by DNP or AZD demonstrates that other signals derived from chloroplasts (e.g., from photosynthetic electron transport) are sufficient to modulate nuclear alternative splicing in this tissue.

## Discussion

Our results reveal a central role for TOR kinase in alternative splicing regulation by light in non-photosynthetic tissues (roots) and, to a certain extent, also in photosynthetic tissues (leaves). As shown previously (15), TOR activation in roots is triggered by sugars that feed glycolysis and the mitochondria. Disruption of the mitochondrial proton gradient by DNP abolishes the light/sucrose-triggered changes on alternative splicing in roots (Fig. 3B) while causing a striking inhibition of RPS6 phosphorylation (Fig. 3D). As TOR kinase is the major regulator of mRNA translation in all eukaryotic cells (24), and light/dark transitions are also known to modulate translation in plants (25), we hypothesized that light and sugars activate the translation of a factor, that in turn, modulates nuclear splicing decisions. Interestingly, 5’ UTRs of mRNAs coding for different *Arabidopsis* SR protein family members (26) are enriched in motifs that provide translational responsiveness to the signaling pathways under consideration (Suppl. Fig. 11, Suppl. Tables 1 and 2). They include a TAGGGTTT motif commonly found in the 5’ UTRs of transcripts, which are rapidly translated after light exposure (25), and a pyrimidine-rich element present in mRNAs that are translationally regulated by TOR (22, 27) (Suppl. Fig. 11, Suppl. Tables 1 and 2).

In terms of alternative splicing regulation, we concluded that at least one main pathway of light modulation involves sugars and TOR kinase. Light is initially sensed by the chloroplast that triggers a signaling pathway that, in turn, involves the mitochondria and TOR in roots. In leaves, even though a similar mechanism is also active, there are other retrograde signals dependent on the photosynthetic electron transport (DCMU sensitive) that can modulate nuclear splicing decisions. Different signaling mechanisms could converge in the activation of translation in the cytosol. Interestingly, it was recently shown that TOR and RPS6 transmit light signals that enhance translation in deetiolating seedlings (28). The participation of a newly translated protein (i.e., a splicing regulator) that moves to the nucleus could be the missing piece communicating different organelles, and signaling pathways, in the cells and explaining underlying retrograde signaling mechanisms (Fig. 3G). The conservation of TOR kinase and its main functions in all eukaryotic cells, together with the involvement of mitochondria, renders the existence of the very same mechanisms of regulation by nutrients in animal cells an intriguing prospect.

## Supporting information

## Acknowledgments

We thank A.R. Kornblihtt, R. Hausler, J. Sheen, A. Köhler, M. Muñoz, P. Duque for materials, discussions and advice. We also thank the “307”, Kornblihtt, Srebrow, Barta and Kalyna groups for the great working atmospheres, A.E. Cambindo Botto, D. Reifer, Viole and Mbututu for extensive and exciting discussions, the IFIBYNE technicians for their support, and L. Giono for her awesome art. This work was supported by the Austrian Science Fund FWF (P26333 to M.K.; DK W1207 and SFBF43-P10 to A.B.) and the Agencia Nacional de Promoción de Ciencia y Tecnología of Argentina to E.P. (PICT 2016 4366). E.P. was an EMBO long term fellow (ALTF_1337-2012), a Marie Curie postdoctoral fellow P330888 (ASRNAbidoPhys) and is now a career investigator from the Consejo Nacional de Investigaciones Científicas y Técnicas of Argentina.

## Author Contributions

E.P. designed and performed the majority of the experiments with the assistance of S.R., L.S., A.F., P.V., M.G.K. and M.A.G.H.. A.F. and A.S. generated RS31-GFP plants and conducted microscopy experiments. C.S, J.W.S.B., C.M, M.K. and A.B. provided input and material to shape the research and the manuscript. M.K., A.B. and E.P. discussed the results and wrote the manuscript with feedback from all the authors.

## Experimental Procedures

### Plant material and growth conditions

For most experiments the *Arabidopsis thaliana* Col-0 ecotype was used as wild type. Transgenic lines used in this study were the RNAi line of TOR (target of rapamycin) 35-7 (20) and transgenic plants expressing At-RS31 protein fused to GFP at the C-terminus. The latter were obtained by transforming wild type plants with constructs that drive the genomic DNA of At-RS31 by either 35S CaMV or its own promoter (about 1 kb upstream of the transcription start site). A detailed description will be published elsewhere. Seeds were stratified for three days in the dark at 4° C and then germinated on Murashige and Skoog medium buffered with 2-(N-Morpholino)ethanesulfonic acid (MS-MES) and containing 1.5 % agar. Plants were grown at a constant temperature of 23 °C under fluorescent lamps emitting white light of an intensity of irradiance between 70 and 100 µmol/m^2^ sec. Growth conditions different from these are indicated in text and figure legends.

### Confocal microscopy

*A. thaliana* plant tissues were mounted in 50% glycerol and analyzed via a Zeiss LSM700 laser scanning confocal microscope using EC Plan-Neofluar 10x/0.3, Plan-Apochromat 40x/1.3 Oil, Plan-Apochromat 63/1.4 Oil in dual-track channel mode or as indicated in the figure legend. GFP and chloroplast autofluorescence were excited with 488 nm laser line, and emissions were recorded at 505 – 530 nm and 650 – 710 nm, respectively. Images were unmixed *in silico* using previously recorded, background-free spectra of chloroplast auto-fluorescence and of GFP.

### Drugs, sugar and low oxygen treatments

Subsequently to growing the plants for two weeks in constant light, they were incubated for 48 hours in the dark. For the treatments with different compounds, plants on agar plates were flooded with 20 mL of liquid MS-MES medium supplemented with the drug or ethanol/dimethyl sulfoxide (vehicle) as a control. This was done after 47 hours of dark treatment (1 hour before the end of the incubation in the dark). Vacuum was applied for five minutes to facilitate drug uptake by the different tissues. The used drugs and their final concentrations were: 15 µM DCMU (3-(3,4-dichlophenyl)-1,1-dimethylurea; Sigma); 20 µM 2,4-DNP (2,4-dinitriphenol; Sigma); and 20 µM AZD-8055 (Chemdea LLC). After the 47 + 1 hours of darkness (with/without drugs) the plants were incubated in light or dark for additional 4 hours. Sucrose was added alone or together with indicated drugs at the specified concentrations (see figure legends). Sorbitol was used as an osmotic control. Sucrose and sorbitol solutions were directly poured onto the agar plates to flood the plants after 47 hours of darkness, letting the plants passively take the sugars for one hour in the dark, or followed by the application of vacuum to facilitate the different compounds uptake by all tissues. After the end of the 48 hours of incubation in the dark, the plants were transferred to light and further incubated for four hours. Controls were kept in the dark.

Low oxygen treatments were carried out in similarly but using isolated (detached) roots instead of seedlings. Briefly, after the 47 hours of darkness, plants were dissected, and isolated roots were treated with sorbitol or sucrose (50 mM). After the 5 minutes of vacuum, the atmosphere of the desiccation chamber used for vacuum infiltration was replaced with air saturated in nitrogen (from a liquid nitrogen canister). The incubation lasted four hours. Air exchange in the desiccation chamber was kept during the whole incubation ensuring that only nitrogen saturated air was able to go in. For the control (normal oxygen), ambient air was used instead of nitrogen saturated air.

### RT- PCR for alternative splicing assessment

Extraction of plant total RNA was carried out using PeqGold (PeqLab) following the manufacturer’s instructions. For cDNA synthesis, 1 µg of RNA was used with the Reverse Transcription System (Promega) and oligo-dT as primer following the manufacturer’s instructions. PCR amplification was performed with Phusion (Thermo) using 2 µl of 1/5 diluted cDNA. The PCR program was: 1) 95º C × 3’, 2) 28-32 cycles of 95º C × 30'', 58-60º C × 30'', 72º C 1’, 3) 72º C × 5’. RT-PCR products were electrophoresed and detected using RedSafe (iNtRON Biotechnology) dye and UV light. The relative intensities of the bands were measured by densitometry using ImageJ software. In particular cases, radioactive alpha-[32P]-dCTP was used in order to assess the alternative splicing changes as described before (7). Results were also validated using Real Time RT-qPCR for individual isoforms and also compared to the results from a high-resolution RT-PCR panel (29). See Supplementary Table 3 for primer sequences.

### Western blot

The organic phase of the PeqGold RNA isolation was used, and proteins were precipitated by addition of cold acetone and subsequent overnight incubation at −20º C. After centrifugation and ethanol washes of the protein pellets 100 µl of Laemmli’s buffer were added. Ten µl of resuspended protein extract were loaded on SDS-PAGE gels. Primary antibodies were RPS6-Phosphorylated (1:5.000) (22), RPS6 total (was obtained similarly to RPS6-P but using an unphosphorylated epitope (22)), anti-histone 3 polyclonal (1:5.000, Agrisera) and anti-cFBP (cytosolic fructose-1,6-bisphosphatase) as loading controls. HRP-conjugated goat anti-rabbit was used as secondary antibody (1:10.000).

### Motif Analysis

The 5’ UTR sequences of the SR genes were obtained from The Arabidopsis Information Resource (ftp://ftp.arabidopsis.org/home/tair/Sequences/blast_datasets/TAIR10_blastsets/). The motif identification was performed using MEME software suite (30) version 4.12.0 with the minimum and maximum motif width set to 6 and 50 nucleotides, respectively, and allowing for reverse complement matches. FIMO tool (31) was used to identify all occurrences of the motifs in the 5’ UTR sequences of the SR genes summarized in supplementary tables (Suppl. Tables 1 and 2). Motifs enriched in the 5’UTRs of light and TOR regulated transcripts were extracted from bibliography (22, 25, 27).

### Statistics

Statistical analyses were carried out using InfoStat (http://www.infostat.com.ar/) 2018e. Same letters indicate means that are not statistically different (p>0.05) for variance analyses with comparisons using Fisher LSD (Least Significant Difference) test from this package.

## Supplementary Materials

Supplementary Figures 1-11 and Supplementary Tables 1-3: see “Suppl. Figs. and Text” file.

