## Supplementary material for "Light remote control of alternative splicing in roots through TOR kinase"

### Suppl. Fig. 1

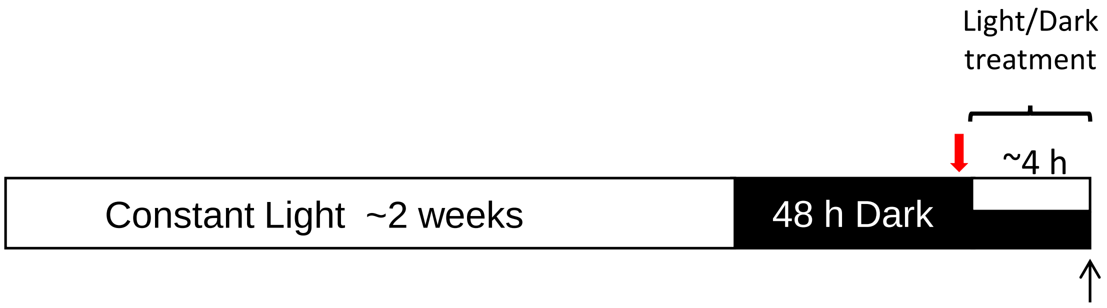

**Suppl. Fig. 1. Standard treatment protocol.** Seedlings were grown in Murashige and Skoog/2-(N-morpholino)ethanesulfonic acid buffered (MS-MES) agar plates (~15 seedlings per 10 cm Ø plate) for a period of two weeks and then transferred for 48 hours to darkness. After this period, seedlings were either transferred to light or kept in darkness for, approximately, additional four hours (light/dark treatment). Sorbitol, sucrose and the used drugs, were added in liquid media (20 mL) on top of the agar-growth media (20 mL) and vacuum was applied in order to facilitate compounds uptake by the different plant tissues. These compounds were added one hour before the end of the 48 hours darkness period (red arrow). Sample collection was performed at the end of the light/dark treatment and it is shown by the black arrow. When needed, prior sample collection, plants were dissected and leaves and roots were collected separatedly (post-treatment dissection). In particular experiments, plants were dissected before light/dark treatments (pre-treatment dissection).

### Suppl. Fig. 2

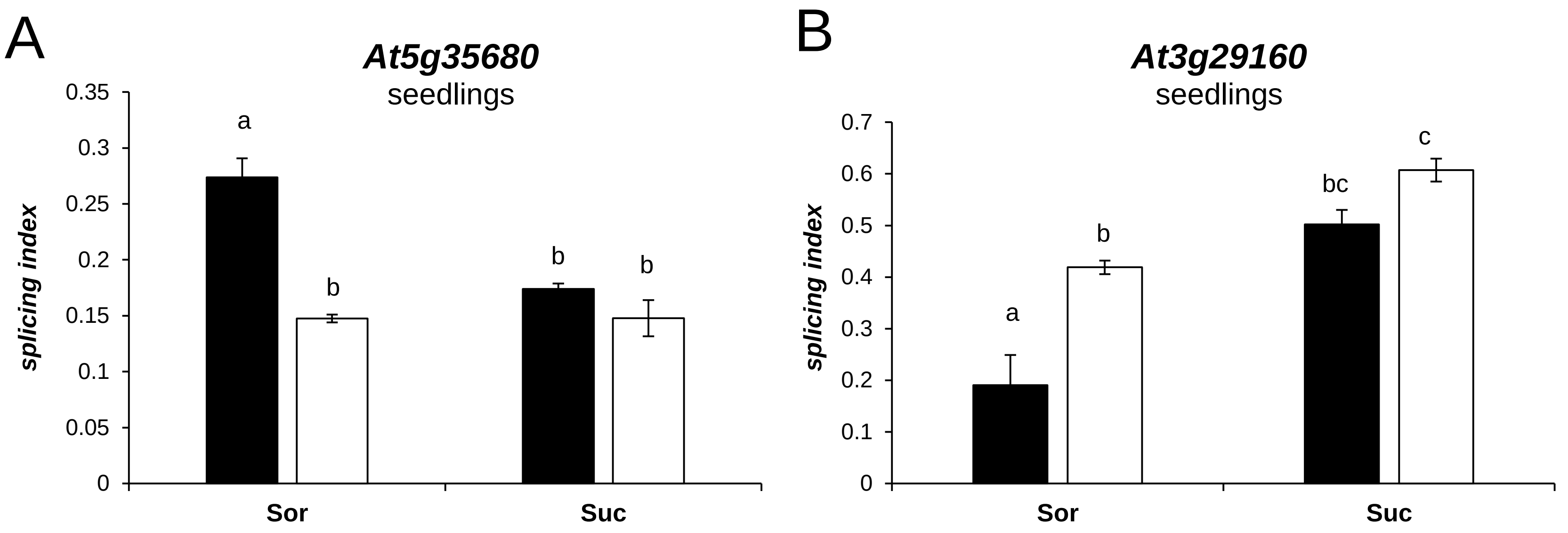

**Suppl. Fig. 2. Sucrose mimics light effects on the regulation of alternative splicing of particular genes. A) AT5G35680, Translation initiation factor 1A (eIF-1A). B) AT3G29160, SNF1-RELATED PROTEIN KINASE 1.2, SNRK1.2.** *A. thaliana* seedlings were grown on MS-MES agar plates (~15 seeds per plate) for a period of 2 weeks under constant light and then incubated in light (white bars) or dark (black bars) for two days. Sorbitol (Sor), or sucrose (Suc) at a 200 mM concentration were added in the plant growth media. The graphs show splicing index (ratio of the longer transcript isoform relative to all the isoforms of a given gene) means ± standard error (n=3). Same letters indicate means that are not statistically different ($p>0.05$). Variance analysis was carried out using the software InfoStat (2018e). Comparisons were made with Fisher LSD (Least Significant Difference) test from this package.

### Suppl. Fig. 3

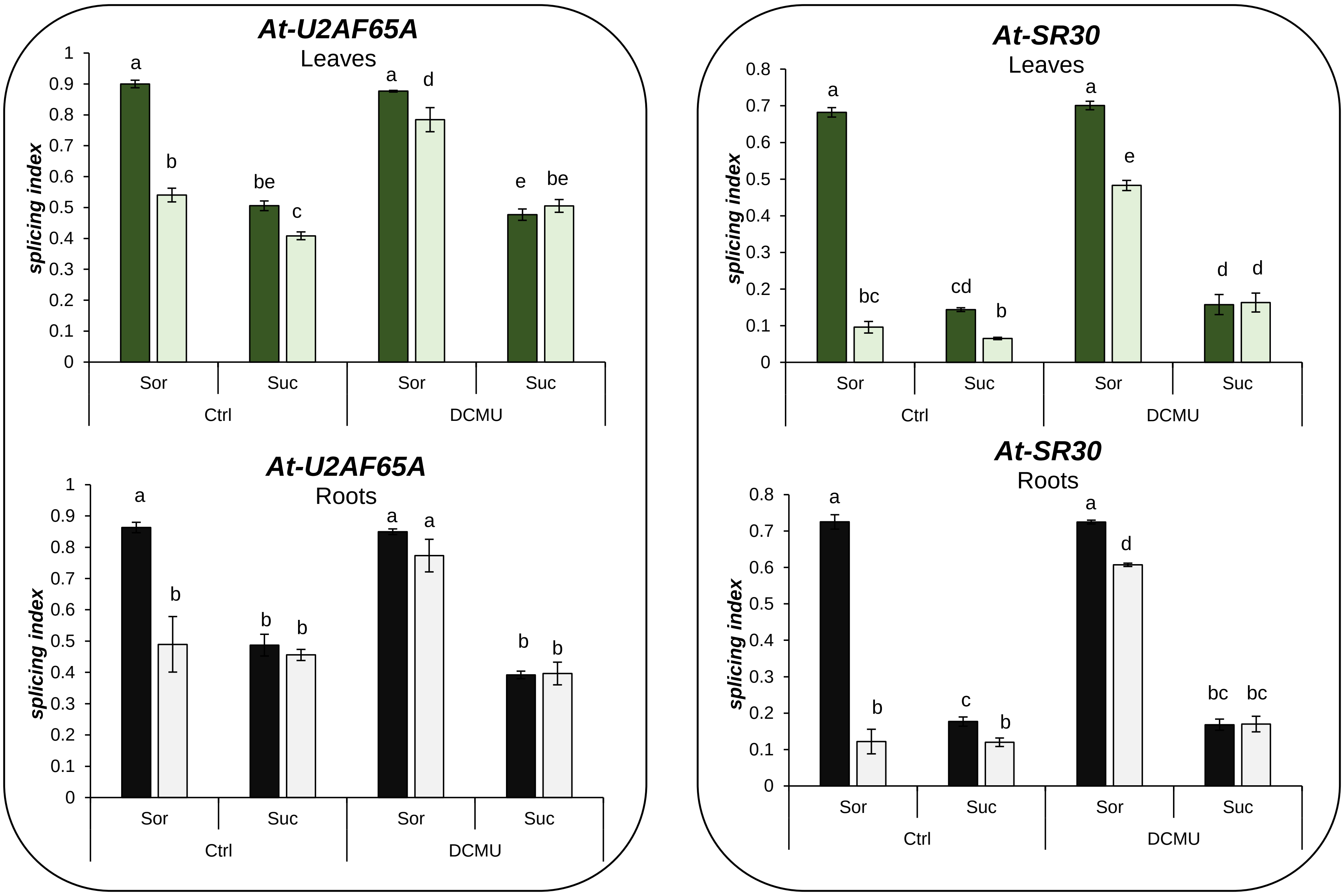

**Suppl. Fig. 3. Sucrose effects on alternative splicing do not involve chloroplasts.** Alternative splicing changes are shown for *At-U2AF65A* (left) and *At-SR30* (right). *A. thaliana* seedlings were grown on MS-MES agar plates (~15 seeds per plate) for a period of two weeks under constant light, then transferred to darkness for 48 hours. Sorbitol (Sor, 100 mM) or sucrose (Suc, 100 mM) supplemented liquid media, with DCMU or without it (ethanol was used as control, Ctrl), were added on top of agar media one hour before the end of the 48 hours darkness period. Vacuum infiltration was applied for five minutes to increase the uptake of the different compounds by all the tissues. After the light (lighter bars) / dark (darker bars) treatments (~4h), leaves (green bars) and roots (grey bars) were dissected for sample collection. The graphs show splicing index means ± standard error (n=4). Same letters indicate means that are not statistically different (p>0.05). Statistics were done using InfoStat with Fisher LSD for comparisons.

### Suppl. Fig. 4

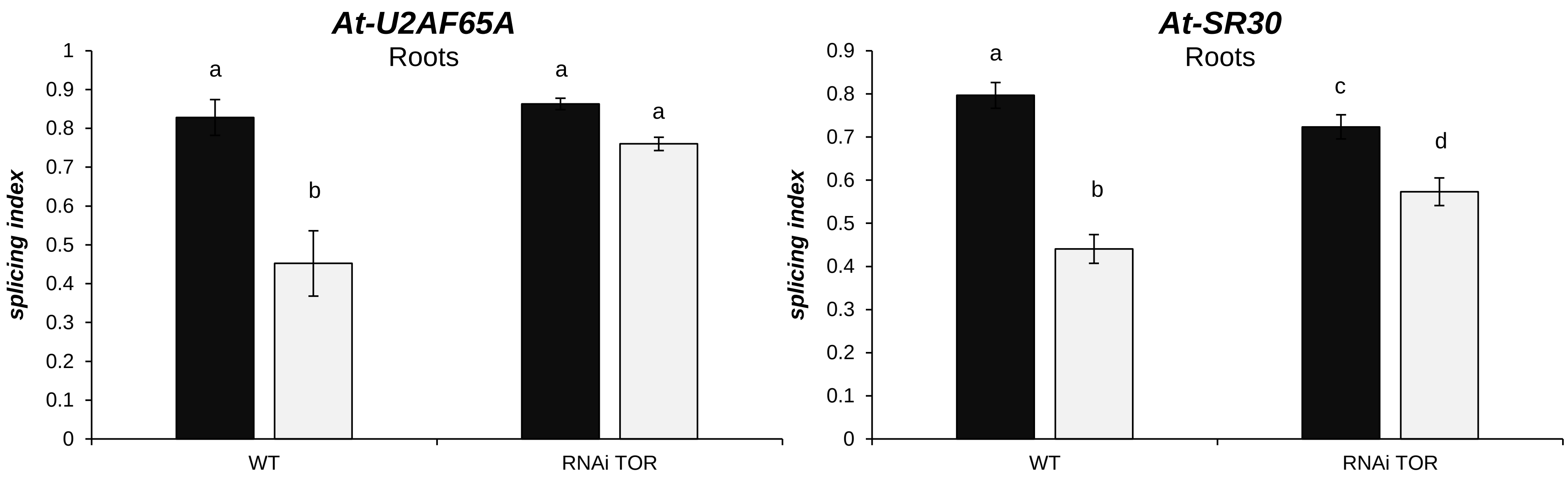

**Suppl. Fig. 4. A transgenic RNAi knockdown line of TOR (RNAi TOR) shows disrupted responses to light on the splicing regulation in roots.** Alternative splicing changes are shown for *At-U2AF65A* and *At-SR30. A. thaliana* wild type (Col-8) and transgenic (35-7 TOR RNAi) line seedlings were grown on MS-MES agar plates (~15 seeds per plate) for a period of two weeks under constant light, then transferred to darkness for 48 hours. After light (lighter bars) / dark (darker bars) treatments for additional four hours, roots were dissected for sample collection. WT, Col-8. The graphs show splicing index means ± standard error (n=4). Same letters indicate means that are not statistically different (p>0.05). Statistical analyses were done using InfoStat with Fisher LSD for comparisons.

### Suppl. Fig. 5

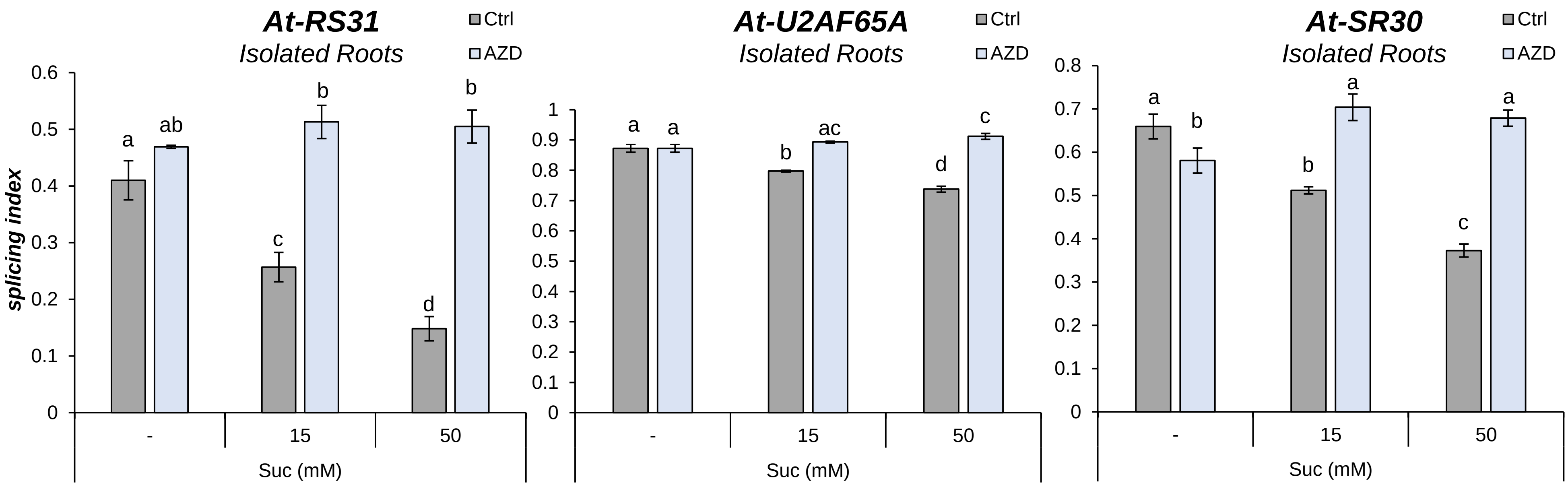

**Suppl. Fig. 5. TOR kinase activity is necessary for isolated (detached) roots to change the alternative splicing of *At-RS31, At-U2AF65A* and *At-SR30* in response to sucrose.** Sucrose (Suc) addition mimics light effects on alternative splicing, reducing the splicing index values in isolated roots in a dose dependent manner. The inhibition of TOR kinase activity by AZD-8055 (AZD) abolishes the effect of sucrose. *A. thaliana* plants were grown on MS-MES agar plates (~15 plants per plate) for a period of two weeks under constant light and then incubated in the dark for 48 hours. Roots were detached and transferred to 6-well plates with liquid media supplemented with sucrose 0, 15 or 50 mM. Sorbitol was used as osmotic control (to reach a total 50 mM concentration, together with sucrose, in every treatment). Twenty µM AZD was used for treatments and dimethyl sulfoxide was used as control (Ctrl). Vacuum was applied for five minutes to increase the uptake of the different compounds. The graphs show splicing index means ± standard error (n=3). Same letters indicate means that are not statistically different ($p>0.05$). Statistical analyses were done using InfoStat with Fisher LSD for comparisons.

Suppl. Fig. 6

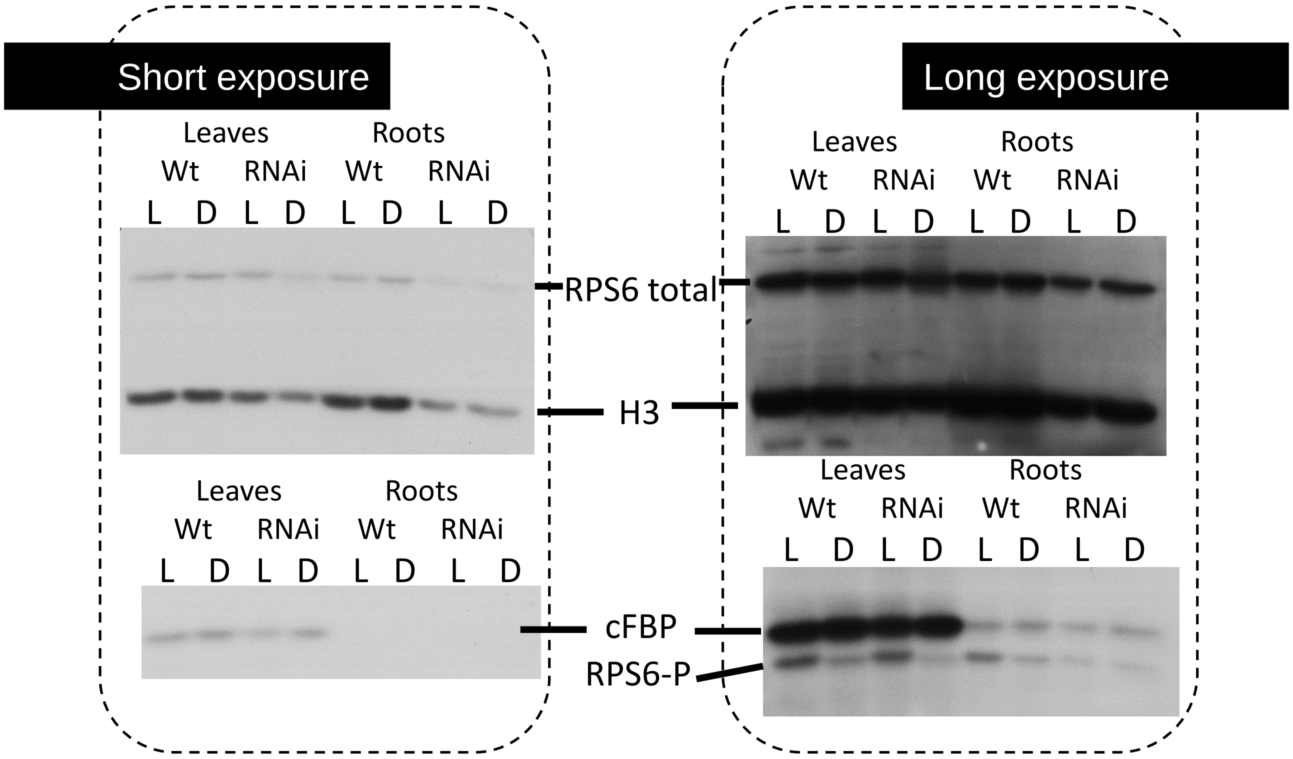

**Suppl. Fig. 6. TOR kinase activity is reduced in a knockdown TOR RNAi line.** Ribosomal protein S6 (RPS6) levels correlate with histone 3 (H3), suggesting its expression is not changing in the tested conditions. RPS6 phosphorylation (RPS6-P) is increased by light (L) in leaves and roots in comparison to dark (D). Note that the arrangement here is first L and then D. This increment in RPS6-P is inhibited in the TOR RNAi line, which expresses an RNAi to knockdown TOR expression. This inhibition is partial in leaves but it is a complete abolishment in roots, mirroring the effects in alternative splicing of the knockdown transgenic line. Images from typical western blot results are shown. Left panel, short exposure; right panel, longer exposure of the same membrane. Antibodies used were, RPS6-P and RPS6, to check the phosphorylation state of RPS6, and H3 and cFBP (cytosolic fructose-1,6-bisphosphatase) as loading controls. *A. thaliana* plants were grown on MS-MES agar plates (~15 plants per plate) for a period of two weeks in constant light and then incubated in the dark for 48 hours. Light/dark treatment was conducted for additional four hours after the 48 hours darkness period.

Suppl. Fig. 7

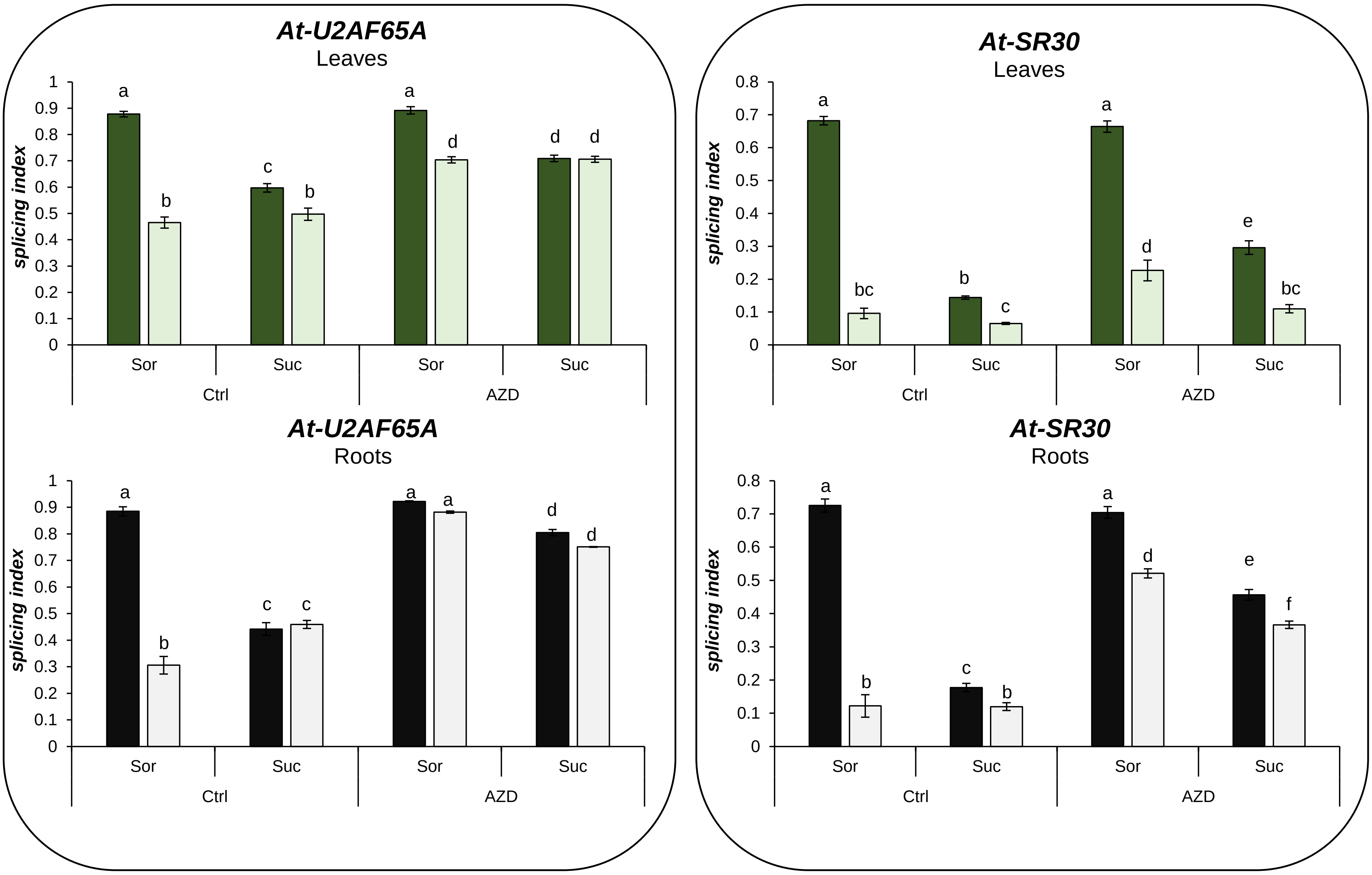

**Suppl. Fig. 7. The inhibition of TOR kinase activity by AZD-8055 disrupts alternative splicing responses.** Alternative splicing changes are shown for *At-U2AF65A* and *At-SR30. A. thaliana* seedlings were grown on MS-MES agar plates (~15 seeds per plate) for a period of two weeks under constant light, then transferred to darkness for 48 hours. Sorbitol (Sor, 100 mM) or sucrose (Suc, 100 mM) supplemented liquid media, with 20 µM AZD-8055 (AZD) or without it (dimethyl sulfoxide as control, Ctrl), were added on top of the agar media one hour before the end of the 48 hours darkness period. Vacuum infiltration was applied for five minutes to increase the uptake of the different compounds by all the tissues. After the light (lighter bars) / dark (darker bars) treatments for additional four hours, leaves and roots were dissected for sample collection. The graphs show splicing index means ± standard error (n=4). Same letters indicate means that are not statistically different (p>0.05). Statistical analyses were done using InfoStat with Fisher LSD for comparisons.

### Suppl. Fig. 8

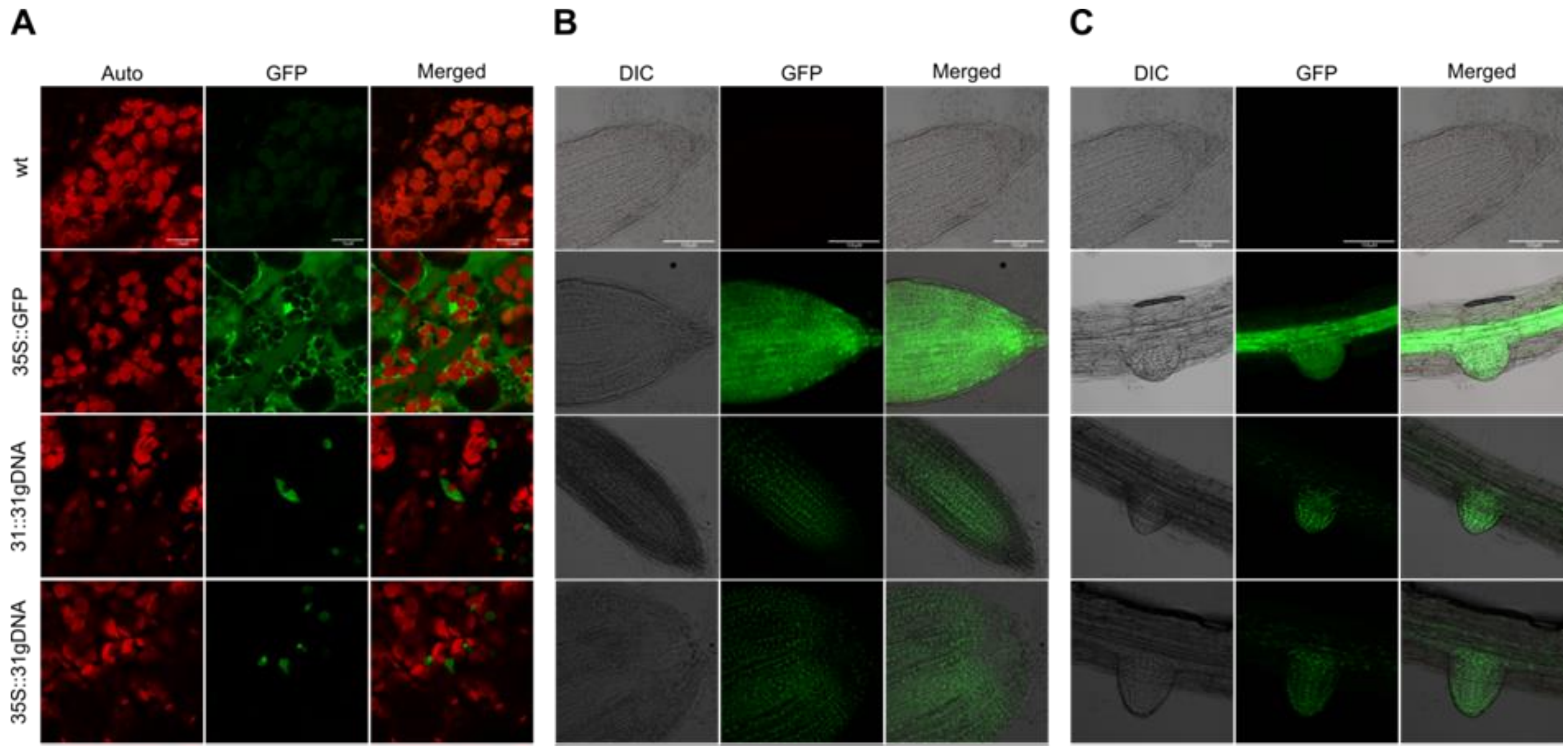

**Suppl. Fig. 8. At-RS31-GFP fusion protein is expressed in cell nuclei of leaves and roots. In root cells the expression of At-RS31-GFP occurs mainly at meristems.** Confocal laser scanning microscopy of leaves **(A)** as well as primary **(B)** and secondary **(C)** roots of living wild type, 35S::GFP and genomic At-RS31-GFP plants under control of either their respective endogenous- or the strong 35S promoter. Wild type and 35S::GFP plants were used as negative and positive controls, respectively. Scale bar represents 10 µm (A) and 100 µm (B,C). For labelling, the –GFP suffix was omitted for At-RS31-GFP transgenic line. Auto, chloroplast autofluorescence. DIC, differential interference contrast.

### Suppl. Fig. 9

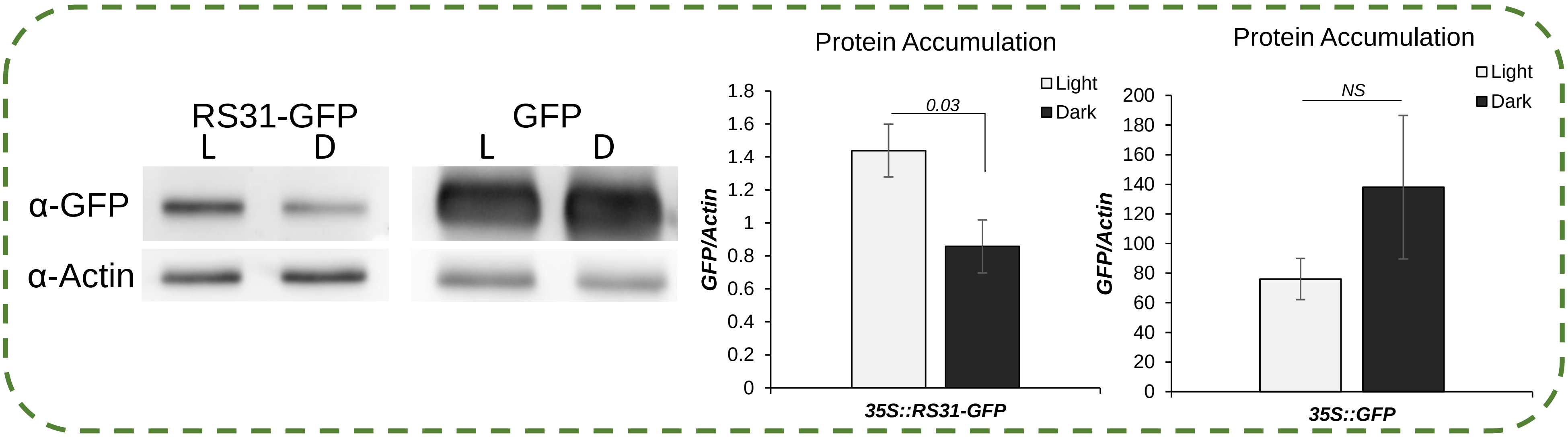

**Suppl. Fig. 9. At-RS31-GFP fusion protein is expressed at higher levels in light than in dark.** Western blots with anti-GFP (Roche) to detect RS31-GFP fusion protein or GFP alone. The transgenic lines express these proteins from a **35S**::**RS31-GFP** or **35S**::**GFP** construct, respectively. Samples were obtained from leaves. Actin (α-Actin) was used as loading control. *A. thaliana* plants were grown on MS-MES agar plates (~15 plants per plate) for a period of two weeks under long day conditions (16:8) and then incubated in the dark for 62 hours. Light (lighter bars) /dark (darker bars) treatment was conducted for additional five hours after darkness. Graphs show protein levels (GFP signal) relative to actin; means ± standard error (n=4). Relevant comparison's p-values (Student's t-test) are shown. *NS*, not significant (p>0.05). Similar results were obtained with seedlings grown in constant light conditions and treated in darkness for two days, also using another antibody (α-GFP; GFP (B-2): sc-9996. Santa Cruz Biotechnology, Inc.).

### Suppl. Fig. 10

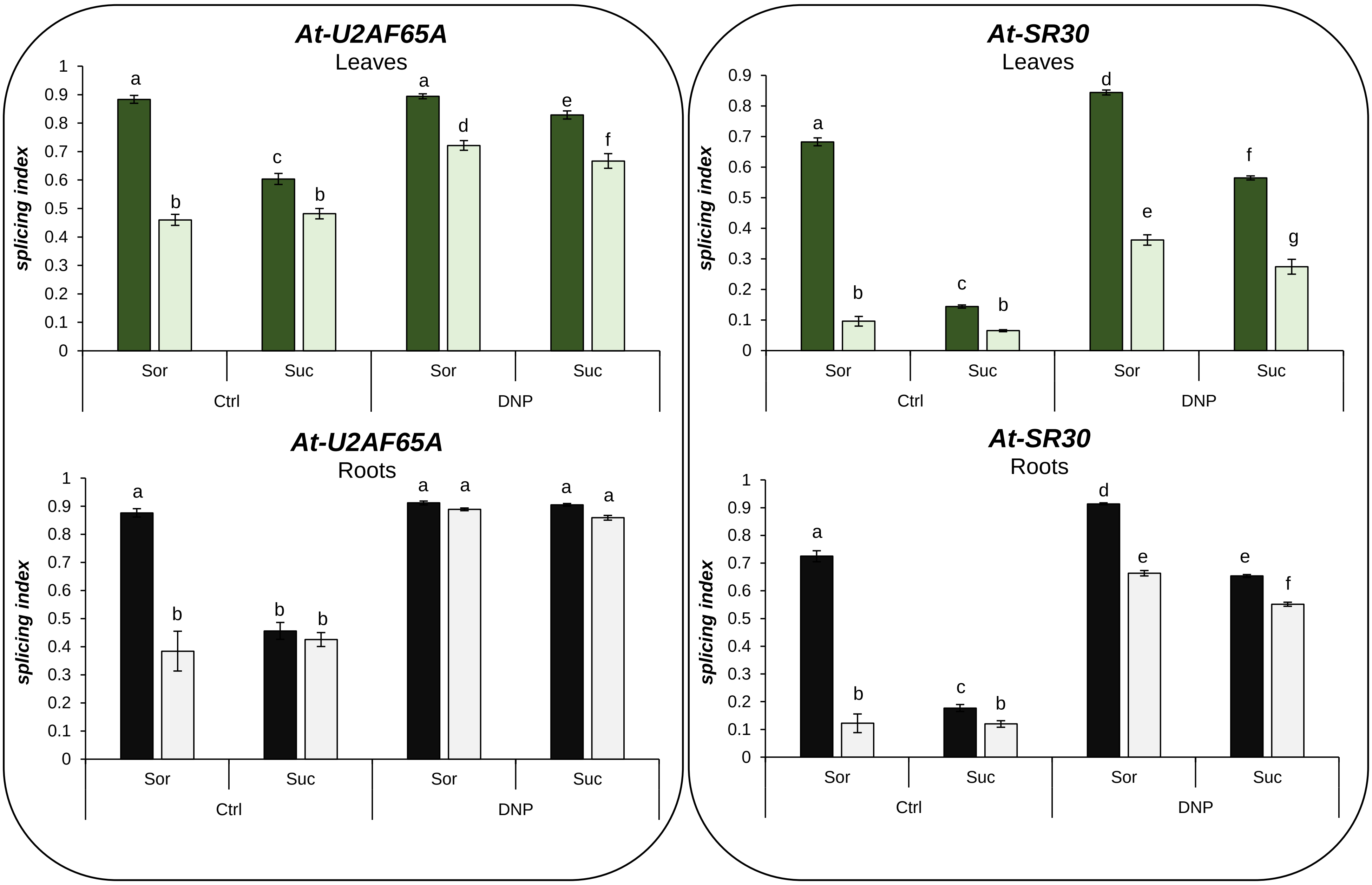

**Suppl. Fig. 10. Proton gradients disruption by an uncoupler obliterates alternative splicing changes induced by light and sucrose in roots.** Alternative splicing changes are shown for *At-U2AF65A* and *At-SR30. A. thaliana* seedlings were grown on MS-MES agar plates (~15 seeds per plate) for a period of two weeks under constant light, then transferred to darkness for 48 hours. Sorbitol (Sor, 100 mM) or sucrose (Suc, 100 mM) supplemented media, with 20 µM DNP (dinitrophenol) or without it (ethanol was used as control, Ctrl), were added on top of the agar media one hour before the end of the 48 hours darkness period. Vacuum infiltration was applied for five minutes to increase the uptake of the different compounds by all the tissues. After light (lighter bars) / dark (darker bars) treatments for additional four hours, leaves and roots were dissected for sample collection. The graphs show splicing index means ± standard error (n=4). Same letters indicate means that are not statistically different (p>0.05). Statistical analyses were done using InfoStat with Fisher LSD for comparisons.

### Suppl. Fig. 11

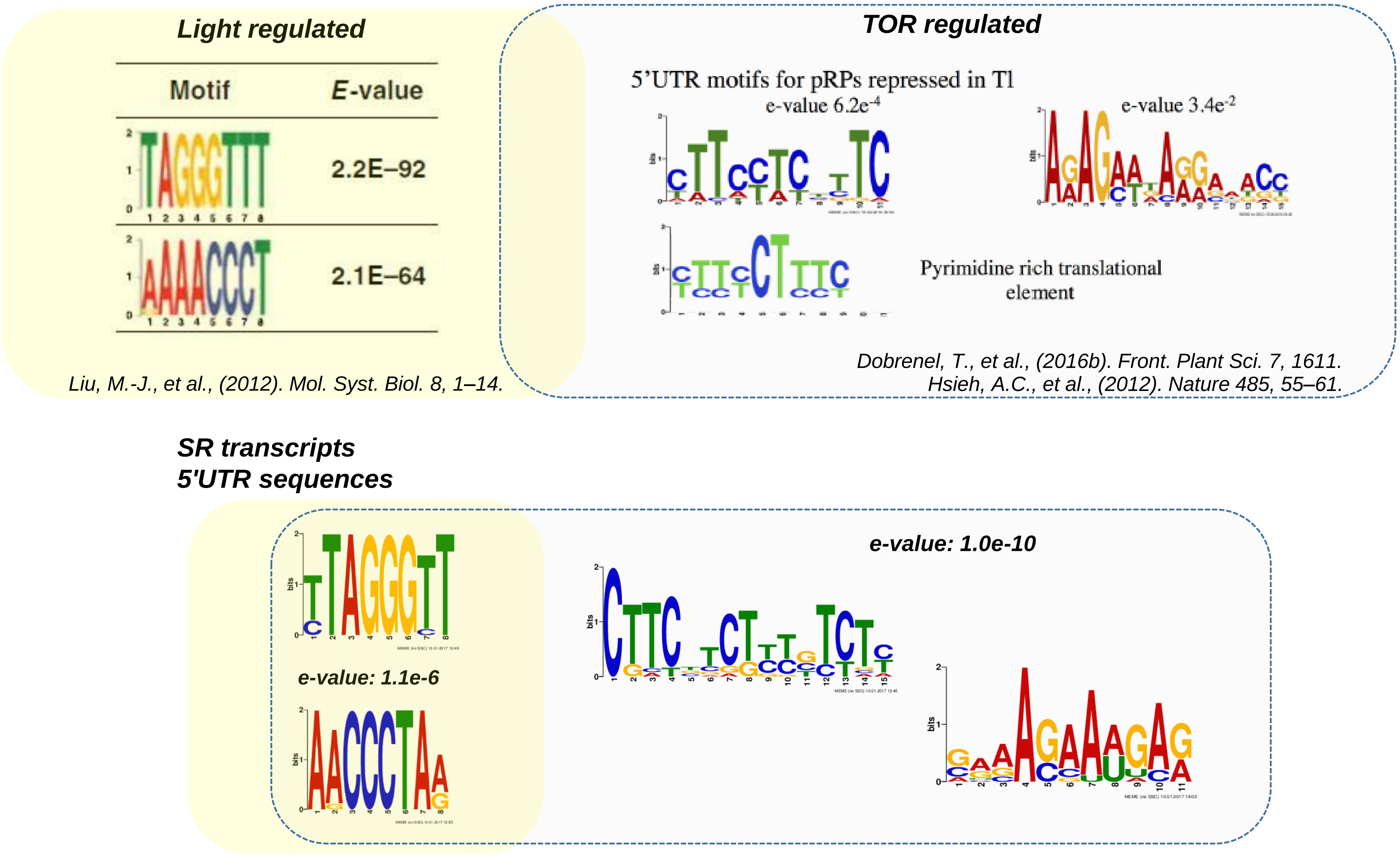

**Suppl. Fig. 11. Over-represented motifs in the 5' UTRs of light and TOR regulated transcripts match with those in the SR protein transcripts.** The 5’ UTR sequences of the SR genes were obtained from The Arabidopsis Information Resource (TAIR10; ftp://ftp.arabidopsis.org/home/tair/Sequences/blast_datasets/TAIR10_blastsets/). The motif identification was performed using the MEME software suite (Bailey and Elkan, 1994; version 4.12.0) with the minimum and maximum motif width set to 6 and 50 nt, respectively, and allowing for reverse complement matches.

### Suppl. Table 1

| Supplementary Table 1: Light-like motif hits* | | | |
|---|---|---|---|
| **A. Mammalian orthologs** | | | |
| **Subfamily** | **Gene/Protein symbol** | **Gene Locus** | **Light-like motif** |
| **SR** ASF/SF2 (SRSF1) orthologs | SR30 | AT1G09140 | – |
| | SR34 | AT1G02840 | AACCCTAA |
| | SR34a | AT3G49430 | AACCCTAG |
| | SR34b | AT4G02430 | – |
| **RSZ** 9G8 (SRSF7) orthologs | RSZ21 | AT1G23860 | AACCCTAA |
| | RSZ22 | AT4G31580 | AACCCTAA |
| | RSZ22a | AT2G24590 | AACCCTAA |
| **SC** SC35 (SRSF2) ortholog | SC35 | AT5G64200 | – |
| **B. Plant specific** | | | |
| **Subfamily** | **Gene/Protein symbol** | **Gene Locus** | **Light-like motif** |
| **SCL** | SCL28 | AT5G18810 | TTAGGGTT |
| | SCL30 | AT3G55460 | TTAGGGTT |
| | SCL30a | AT3G13570 | AACCCTAA |
| | SCL33 | AT1G55310 | – |
| **RS2Z** | RS2Z32 | AT3G53500 | TTAGGGTT |
| | RS2Z33 | AT2G37340 | TTAGGGCT |
| **RS** | RS31 | AT3G61860 | CTAGGGTT |
| | RS31a | AT2G46610 | CTAGGGTT |
| | RS40 | AT4G25500 | CTAGGGCT |
| | RS41 | At5G52040 | CTAGGGTT |

**Suppl. Table 1. Presence and identification of over-represented motifs in the 5' UTRs of SR protein coding transcripts that are enriched in transcripts highly translated upon illumination.** The 5' UTR sequences of the SR genes were obtained from The Arabidopsis Information Resource (TAIR10; ftp://ftp.arabidopsis.org/home/tair/Sequences/blast_datasets/TAIR10_blastsets/). The motif identification was performed using FIMO tool (Grant et al., 2011). Light-like motifs were described in *Liu, M.-J., et al., (2012). Mol. Syst. Biol. 8, 1–14.*

### Suppl. Table 2

| Supplementary Table 2: TOR-like motif hits | | | |
|---|---|---|---|
| **A. Mammalian orthologs** | | | |
| **Subfamily** | **Gene/Protein symbol** | **Gene Locus** | **TOR-like motif** |
| **SR**<br>ASF/SF2 (SRSF1) orthologs | SR30 | AT1G09140 | CATCTTCTTCTTCTT-------<br>--<br>---CTTCTTCTTCTTCTT----<br>--<br>------CTTCTTCTTCTTTCT-<br>--<br>---------<br>CTTCTTCTTTCTCGA |
| | SR34 | AT1G02840 | ACGACCAACAGGAAG |
| | SR34a | AT3G49430 | CGACCTCTCTCTCTC |
| | SR34b | AT4G02430 | CTTCTTCTCTTCCAT |
| **RSZ**<br>9G8 (SRSF7) orthologs | RSZ21 | AT1G23860 | - |
| | RSZ22 | AT4G31580 | AAAACACAGAAAAAG |
|  | RSZ22a | AT2G24590 | ACGAGCAAGAGAAAG-------<br>-<br>--------<br>GAGAAAGAGCCAAAG |
| **SC**<br>SC35 (SRSF2) ortholog | SC35 | AT5G64200 | CTTCGTCTCTCTCTC |
| **B. Plant specific** | | | |
| **Subfamily** | **Gene/Protein symbol** | **Gene Locus** | **TOR-like motif** |
| **SCL** | SCL28 | AT5G18810 | CGATTTCTTCCTCTC------<br>------CTTCCTCTCTCCCTC |
| | SCL30 | AT3G55460 | CTTCAGATCTGTCTT |
| | SCL30a | AT3G13570 | CTTCCCCTGTGTTTT |
| | SCL33 | AT1G55310 | CTTCTCCGTCGTTCC |
| **RS2Z** | RS2Z32 | AT3G53500 | CTCCTTCTTCGTCTC---<br>---CTTCTTCGTCTCTCC |
| | RS2Z33 | AT2G37340 | CTTCCCCTTTCCTTC |
| **RS** | RS31 | AT3G61860 | CGTCGTCGTCGTCTA |
|  | RS31a | AT2G46610 | CGTCGTCGTCGTCTT---<br>---CGTCGTCGTCTTCTA<br><br>CTCCGTCTTCTTCGC---<br>---CGTCTTCTTCGCCGA<br><br>CTTTCTCGTCGTCGT------<br>------CGTCGTCGTCTCTTC |
| | RS40 | AT4G25500 | CTTCTTCTTCTTCTA |
| | RS41 | At5G52040 | GAGAGAGCCTCGAAG |

**Suppl. Table 2. Presence and identification of over-represented motifs in the 5' UTRs of SR protein coding transcripts that are enriched in mRNAs regulated by TOR at the level of translation.** The 5' UTR sequences of the SR genes were obtained from The Arabidopsis Information Resource (TAIR10; ftp://ftp.arabidopsis.org/home/tair/Sequences/blast_datasets/TAIR10_blastsets/). The motif identification was performed using FIMO tool (Grant et al., 2011). TOR-like motifs were described in *Dobrenel, T., et al., (2016b). Front. Plant Sci. 7, 1611* and *Hsieh, A.C., et al., (2012). Nature 485, 55–61.*

### Suppl. Table 3

| Oligos | | |
|---|---|---|
| **Alternative Splicing (RT-PCR)** | **name** | **sequence** |
| At-RS31 | RS31-Fw | cggttgttcgacaagtatg |
| | RS31-Rv | tgtaggcttcagatttgaag |
| | RS31n-Spl-Fw | tcggatctggaacggttg |
| | RS31n-Spl-Rv | cagtgtctttgtaggcttcag |
| At-SR30 | SR30-Fw | gctatacagctctgtctcaag |
| | SR30-E6-Fw | catgcgcaaagctggagatg |
| | SR30-Rv | tttcattttcaaccagatatcac |
| At-U2AF65A | pp198-Fw | ggatgagcttagagatgatgagg |
| | U2AF65-E10-Fw | tgcacagcagcaaatagctt |
| | pp198-Rv | ggcctgccactggctgaccattgg |
| eIF-1A (AT5G35680) | pp48- Fw | cctaatcactttcaacaactc |
| | pp48-Rv | ccgtcaatgcacataacgtc |
| SNRK1.2 (AT3G29160) | pp343- Fw | cctgactcagctctgcgtcacc |
| | pp343- Rv | cccaattccaagagttttacc |
| **Real Time (RT-qPCR)** | **name** | **sequence** |
| At-RS31 mRNA1 | mRNA1-Fw1 | agtgttcgtcggcaatttcg |
| | mRNA1,3-Rv | ctttgcccattcaactgataacc |
| | mRNA1-Fw2 | cgagtggacatgaaatctggata |
| At-RS31 mRNA3 | mRNA3-Fw | tgtgctttcttttgttgcaggata |
| | mRNA3-Rv | ctttgcccattcaactgataacc |
| | mRNA1,3-Rv | ctttgcccattcaactgataacc |
| At-RS31 total expression | RS31-Fw | acactttgagccctatggtaag |
| | RS31-Rv | atcatctctttcatcgtcatctttc |
| At-PP2a (house-keeping) | PP2A-Fw | taacgtggccaaaatgatgc |
| | PP2A-Rv | gttctccacaaccgcttggt |
| HRE1 (At1g72360) | HRE1-Fw | ggcctctgccttatccctctgt |
| | HRE1-Rv | gcgtaaacccgtctcagtgagtg |

**Suppl. Table 3. Sequences and names of the used oligos in the study.**
